# Resistome, Virulome, Mobilome, And Biosynthetic Gene Clusters Adaptations of *Acinetobacter Baumannii* Mexican Strains Across the Pre- and of the COVID-19 Period: Insights from Whole-Genome Sequencing

**DOI:** 10.64898/2026.03.22.712163

**Authors:** Moisés A. Alejo, Miriam Sarahi Lozano Gamboa, Brian Muñoz Gomez, Karla G. Hernández Magro Gil, Rodolfo García-Contreras, Rachel J. Whitaker, Anastasio Palacios Marmolejo, Corina-Diana Ceapă

**Affiliations:** Microbiology Laboratory, Department of Chemistry of Natural Products, Institute of Chemistry, National Autonomous University of Mexico (UNAM), Mexico; Laboratorio Estatal de Salud Pública del Estado de Aguascalientes (LESP), México / Escuela de Medicina Universidad Cuauhtémoc Plantel Aguascalientes; Departamento de Microbiología y Parasitología, Facultad de Medicina, Universidad Nacional Autónoma de México, Ciudad de México, México; Carl R. Woese Institute for Genomic Biology, University of Illinois Urbana-Champaign, Urbana, Illinois, USA; Department of Microbiology, University of Illinois Urbana-Champaign, Urbana, Illinois, USA

**Keywords:** antimicrobial resistance AMR), genomic surveillance, healthcare associated infections (HAI), horizontal gene transfer (HGT), whole genome sequencing (WGS) analysis

## Abstract

**Background:** *Acinetobacter baumannii* is a critical multidrug-resistant pathogen whose genomic landscape in Mexico has been reshaped by the COVID-19 pandemic. While global studies have highlighted distinctive sequence type distributions, systematic analyses in Mexico remain limited.

**Methods:** We analyzed 194 genomes, including 47 newly sequenced post-COVID isolates (MIQ), alongside 147 publicly available genomes (HPG). Whole-genome sequencing was combined with phylogenetic reconstruction, resistome and virulome profiling based on gene presence and absence, mobilome analysis, and biosynthetic gene cluster (BGC) characterization.

**Results:** Two major clades dominated by Oxford STs 758, 208, 417, and 369 were identified. Resistome profiling uncovered 128 distinct resistome profiles (combinations of genes) and 44 emerging antimicrobial resistance genes (ARGs), with an increased number of resistance genes in the strains obtained during the pandemic. Virulome analysis revealed enrichment of metabolic adaptation genes (*argG*, *carA*, *ilvC*) in MIQ strains. Mobilome profiling demonstrated enrichment of ISAbA1 and ISAbA3 elements, known to mobilize carbapenemase genes. BGC analysis showed conserved siderophores involved in virulence, alongside diversification of the secondary metabolite repertoires in MIQ genomes. Additional observations included geographic mixing of clades across Jalisco, Aguascalientes, and Mexico City and referral bias toward carbapenemase-positive isolates.

**Conclusion:** The genomic landscape of *A. baumannii* in Mexico has diversified post-COVID, with evidence of inter-regional transmission, referral bias, virulome expansion, mobilome-driven ARG dissemination, and metabolic adaptation. These findings underscore the urgent need for coordinated genomic surveillance, functional and clinical validation of adaptation signals, and regionally integrated infection control strategies to mitigate resistance trajectories.

**Graphical abstract:** Integrative genomic profiling of *A. baumannii* isolates from Mexico. Workflow summarizing the analysis of 194 genomes (147 historic public genomes, 47 novel MIQ strains). Clinical isolates were identified by MALDI Biotyper and tested with BD Phoenix™ M50. Whole genome sequencing enabled phylogenetic and MLST analyses, resistome and virulome profiling, mobilome characterization, and BGC identification using BIGSCAPE and antiSMASH. Outputs include phylogenetic clustering, resistance/virulence gene distributions, and biosynthetic potential across Mexican regions.

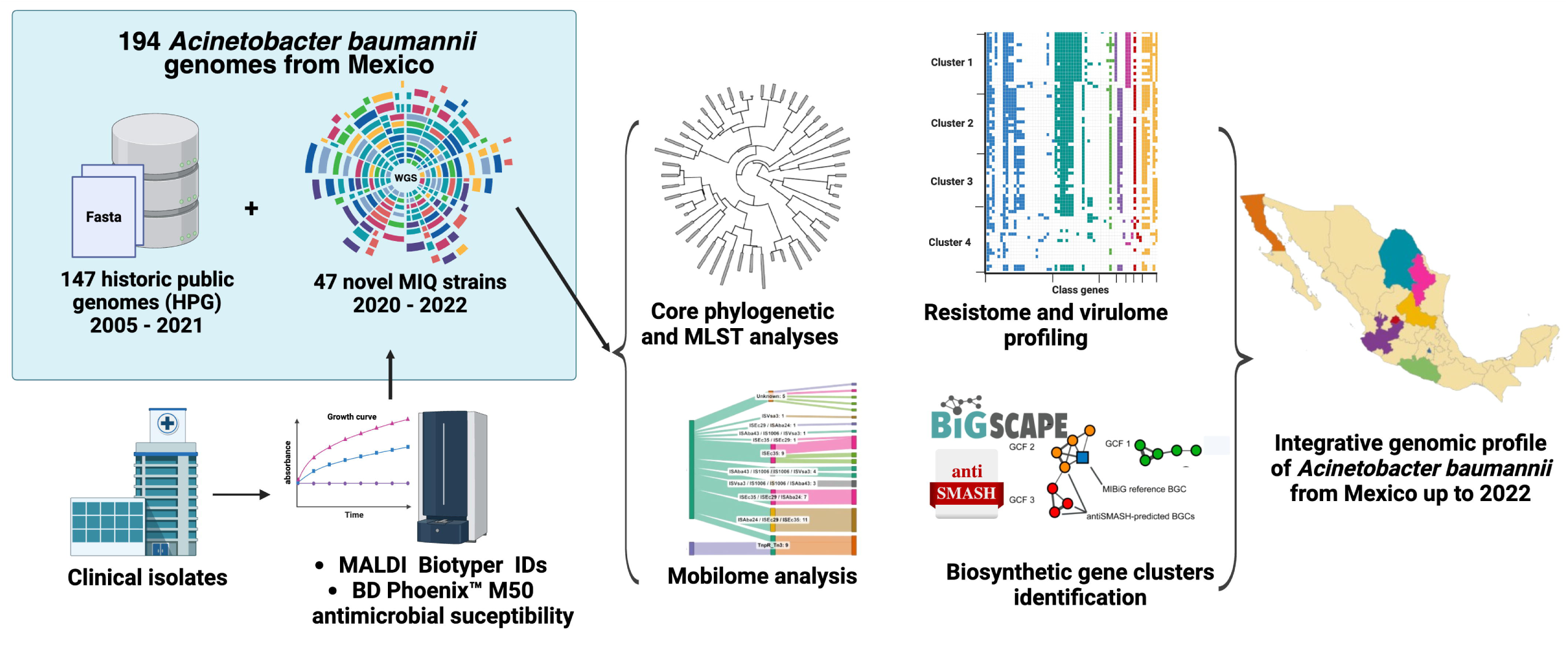

## Introduction

Widespread empirical antibiotic use in COVID-19 patients has led to a significant increase in secondary infections caused by resistant bacteria (Castillo-Bejarano et al., 2023). Among these, *Acinetobacter baumannii* coinfections in ICU settings have surged, frequently exhibiting extensive drug resistance (XDR) or pan-resistance, coupled with high virulence potential, presenting geo-economical specificities (Barrios-Camacho et al., 2025).

Recently, global studies indicated that in the pre-pandemic period, Mexico presented sequence type (ST) patterns that differ from neighboring countries (Fernández-Vázquez et al., 2023). Understanding the genomic traits that underpin these phenotypes is essential for effective infection control and the development of therapeutic strategies.

The persistence and pathogenicity of *A. baumannii* are largely attributed to its remarkable ability to evolve and acquire resistance to multiple drugs through diverse mechanisms, including the production of extended-spectrum β-lactamases (ESBLs), carbapenemases, efflux pumps, and reduced membrane permeability (Gomes Chagas et al., 2024; Manobanda Nata and Jaramillo Ruales, 2023). Beyond the resistome, the virulome encompasses genes responsible for adherence, biofilm formation, and immune evasion, which contribute to the bacterium’s virulence. The mobilome, consisting of plasmids, transposons, and integrative conjugative elements, plays a pivotal role in the horizontal gene transfer (HGT) of resistance and virulence determinants (Fernández-Vázquez et al., 2023). Additionally, the specialized metabolome, which includes biosynthetic gene clusters (BGCs) responsible for specialized metabolites, may influence bacterial fitness, interspecies competition, and interactions with the host and microbiota (Djahanschiri et al., 2022; Djahanschiri et al., 2024).

Collectively, these components highlight the complexity of *A. baumannii* adaptability and underscore the urgent need for comprehensive genomic studies to inform effective countermeasures.

In this study, we conducted a comprehensive genomic investigation using whole-genome sequencing (WGS) and *in silico* analyses of clinical isolates collected from Aguascalientes, Mexico City, and neighboring states in Central Mexico during the COVID-19 pandemic and post-pandemic periods (2020–2023). To provide robust regional and temporal comparisons, we integrated all publicly available genomes characterized across Mexico from the pre-COVID period. This approach provides an integrative and current view of the *A. baumannii* genomic landscape in Mexico, shedding light on the evolution of resistance patterns, virulence factors, mobile genetic elements, and specialized metabolite profiles over time. Our findings aim to support the development of targeted therapies and antimicrobial stewardship programs tailored to the Mexican context.

## Methods

### Ethics and Biosafety

This study was approved by the Promotora Médica Aguascalientes S.A. de C.V. Ethics and Biosafety Committee (Approval Nos. 100.cbpma.2025 and 2937.ceipma.2025), registered with the national regulatory agency COFEPRIS. The protocol encompassed observational, descriptive, and retrospective analyses of multidrug-resistant *Acinetobacter baumannii*, *Escherichia coli*, and *Klebsiella* spp. isolates collected from hospitalized post-COVID-19 patients between 2020 and 2023. Informed consent was obtained in accordance with Mexican regulations (NOM-012-SSA3-2012), and annual progress reports were submitted to ensure compliance with biosafety and clinical research standards. All procedures adhered to WHO guidelines for antimicrobial resistance surveillance and international ethical frameworks for genomic data sharing (PAHO/WHO, 2023).

### Clinical Samples

Between March 2021 and December 2023, clinical specimens were collected from hospitalized patients at major hospitals in Aguascalientes, including Tercer Milenio General Hospital, General Hospital of Zone No. 1 and No. 2, General Hospital of Pabellón de Arteaga, General Hospital of Rincón de Romos, General Hospital of Calvillo, Women’s Hospital of Aguascalientes, and Centenario Miguel Hidalgo Hospital. Additional isolates were obtained from the Faculty of Medicine, UNAM, and forwarded to the State Public Health Laboratory of Aguascalientes (LESPA).

Isolates were transported under approved biosafety protocols, including triple packaging, cold chain maintenance at 4 °C, and delivery within 48 hours. Species-level identification was performed using MALDI-TOF MS (Prieto et al., 2024). Antimicrobial susceptibility testing (AST) was conducted using the BD Phoenix™ M50 system, with interpretation based on CLSI guidelines (CLSI M100, 2023). Extended-spectrum β-lactamase (ESBL) and carbapenemase production were confirmed using phenotypic panels. Because LESPA receives isolates referred for confirmation by medical practitioners, the dataset is enriched for resistant or clinically severe cases, likely representing a local high-risk subset of hospital circulating pathogens (Castillo-Bejarano et al., 2023).

### Selection of strains for sequencing

Strains were randomly selected from clinical isolates submitted to the LESP, most of which were multidrug-resistant and pan-resistant. The collection reflects this natural bias toward high-risk cases, and final inclusion depended solely on staff time and reagent availability, without any intentional bias toward additional characteristics.

### Whole Genome Sequencing

Forty-seven clinical strains from patients with lung disease in Central Mexico were cultured on Columbia agar and confirmed by MALDI Biotyper®. Genomic DNA was extracted using the NEB Monarch® kit according to manufacturer’s instructions. Libraries were prepared with the Illumina MiSeq™ DNA Prep kit v3 and sequenced on the Illumina MiniSeq platform using paired-end 2×150 bp reads.

Comparative genomic analyses of *Acinetobacter baumannii* strains were conducted using a standardized bioinformatics workflow designed to ensure reproducibility and robustness. Raw sequencing reads were first subjected to quality control with FastQC v0.11.9, followed by adapter and low-quality base trimming using Trimmomatic v0.39. High-quality reads were assembled de novo with SPAdes v3.9.0, and assemblies were annotated using Prokka v1.14.6 and the RAST toolkit implemented in the BV-BRC v3.30.19 platform.

### Data Availability

All genomes generated in this study are deposited under BioProject PRJNA1291376 at NCBI. Associated metadata—including collection sites, dates, and sources—are provided in Supplementary Table 1. Draft assemblies available in NCBI were annotated using the NCBI Prokaryotic Genome Annotation Pipeline, while data presented in the text were annotated with SPAdes. Strains are preserved in the Microbiology Laboratory Pathogen Strain Collection, Institute of Chemistry, UNAM, and are available upon request to qualified researchers, especially in support of much needed projects addressing antimicrobial resistance in Latin America (Gomes Chagas et al., 2024; Manobanda Nata and Jaramillo Ruales, 2023).

### Phylogenetic Analysis

Phylogenetic reconstruction was performed using a combined dataset of newly sequenced MIQ isolates and historic published genomes (HPG) retrieved from GenBank (Supplementary Table 1), thus forming a complete HPG-MIQ dataset used for the following analyses.

Core genome alignments were generated using Roary v3.13.0, and phylogenetic trees were reconstructed with IQ-TREE v2.1.2 under the maximum likelihood framework with 1,000 bootstrap replicates. Assemblies were uniformly annotated with Prokka v1.14.6, and STs were assigned using the PubMLST database. Metadata (hospital, state, year, Pasteur and Oxford ST schemes) were incorporated into the tree visualization in iTOL v6, allowing contextual interpretation of lineage distribution across clinical and environmental sources.

### MLST analysis

*In silico* multilocus sequence type (MLST) profiles were assigned using the sequences for MLST markers for the Oxford typing scheme (*gltA*, *recA*, *cpn60*, *gyrB*, *gdhB*, *rpoD*, and *gpi*) and Pasteur typing scheme (*gltA*, *recA*, *cpn60*, *fusA*, *pyrG*, *rpoB*, and *rplB*). Sequences were downloaded from the pubMLST database (http://pubmlst.org/abaumannii) and then used to extract allele sequences from each of the *A. baumannii* genomes using the BLASTN tool. The extracted sequences were then uploaded to the pubMLST database to assign both existing and novel sequence types.

### Resistome and Virulome Analysis

For resistome profiling, raw assemblies were screened with the BV-BRC K-mer pipeline, complemented with BLAST searches against the Comprehensive Antibiotic Resistance Database (CARD) and the National Database of Antibiotic Resistant Organisms (NDARO). In addition, we applied SraX, a pipeline optimized for CARD-based annotation. Following annotation, redundant entries were removed and homologous genes were grouped to generate a non-redundant resistome dataset suitable for comparative analyses.

Resistome diversity was quantified using Bray–Curtis dissimilarity indices, which measure compositional differences in gene content between isolates. The resulting distance matrices were visualized through Principal Coordinates Analysis (PCoA) to identify clustering patterns. To further assess group separation, k-means clustering was applied, and statistical robustness was validated using PERMANOVA (permutational multivariate analysis of variance).

### Mobilome Analysis

Mobile genetic elements (MGEs) were systematically characterized for each genome using VRprofile2. For each isolate, we quantified the number and diversity of MGEs. Descriptive statistics were applied to compare mobilome composition between pre-COVID and post-COVID isolates, including median counts per genome and prevalence of specific replicon or transposon families.

To visualize mobilome dynamics, we employed SankeyMATIC to generate Sankey diagrams illustrating gene flow between chromosomes, plasmids, and MGEs.

### Specialized metabolism

All genomes were analyzed with antiSMASH v7.0 (intermediate exhaustiveness). Biosynthetic gene clusters (BGCs) were compared descriptively. Similarity networks of BGCs were constructed with BiG-SCAPE v2.0 and visualized with Cytoscape, referencing MIBiG v3.1 characterized clusters.

## Results

### Phylogenetic Landscape

The HPG set included *A. baumannii* strains collected between 2011 and 2018 from diverse Mexican hospitals and institutions (e.g., Hospital Civil de Guadalajara, Instituto Nacional de Cancerología, Hospital Infantil de México Federico Gómez, ISSSTE, and Hospital Universitario de Nuevo León), spanning multiple sequence types (ST1, ST2, ST156, ST422, among others). This integration provided temporal and geographic breadth, enabling direct comparison of post-COVID MIQ isolates with pre-COVID epidemic lineages.

Our genomic survey revealed two major clades of *A. baumannii* circulating in Mexico (HPG-MIQ), cutting across geography and time (Figure 1 A and B). The phylogenetic trees show that these dominant lineages are not confined to single states but circulate across Jalisco, Aguascalientes, and Mexico City.

**Figure 1.**
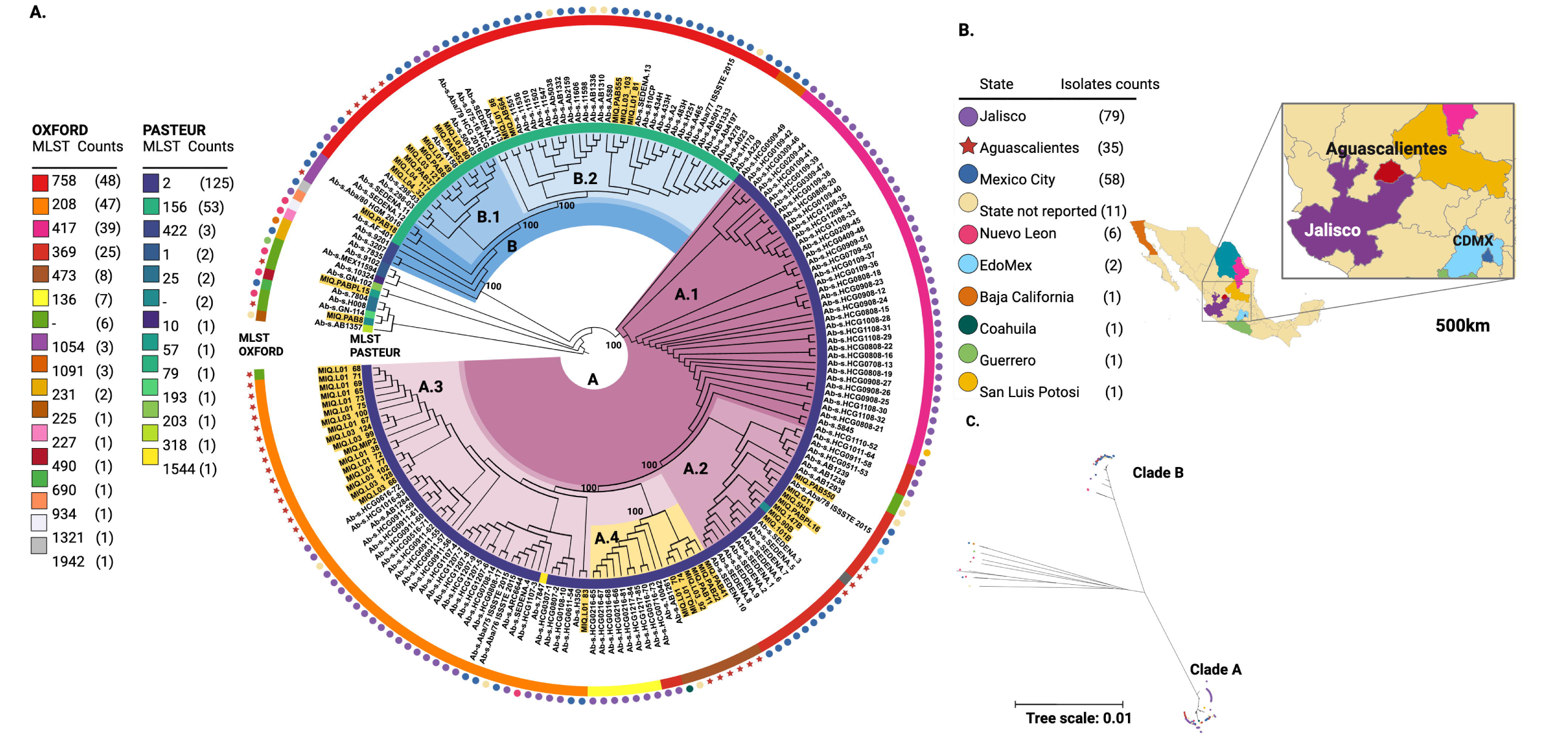
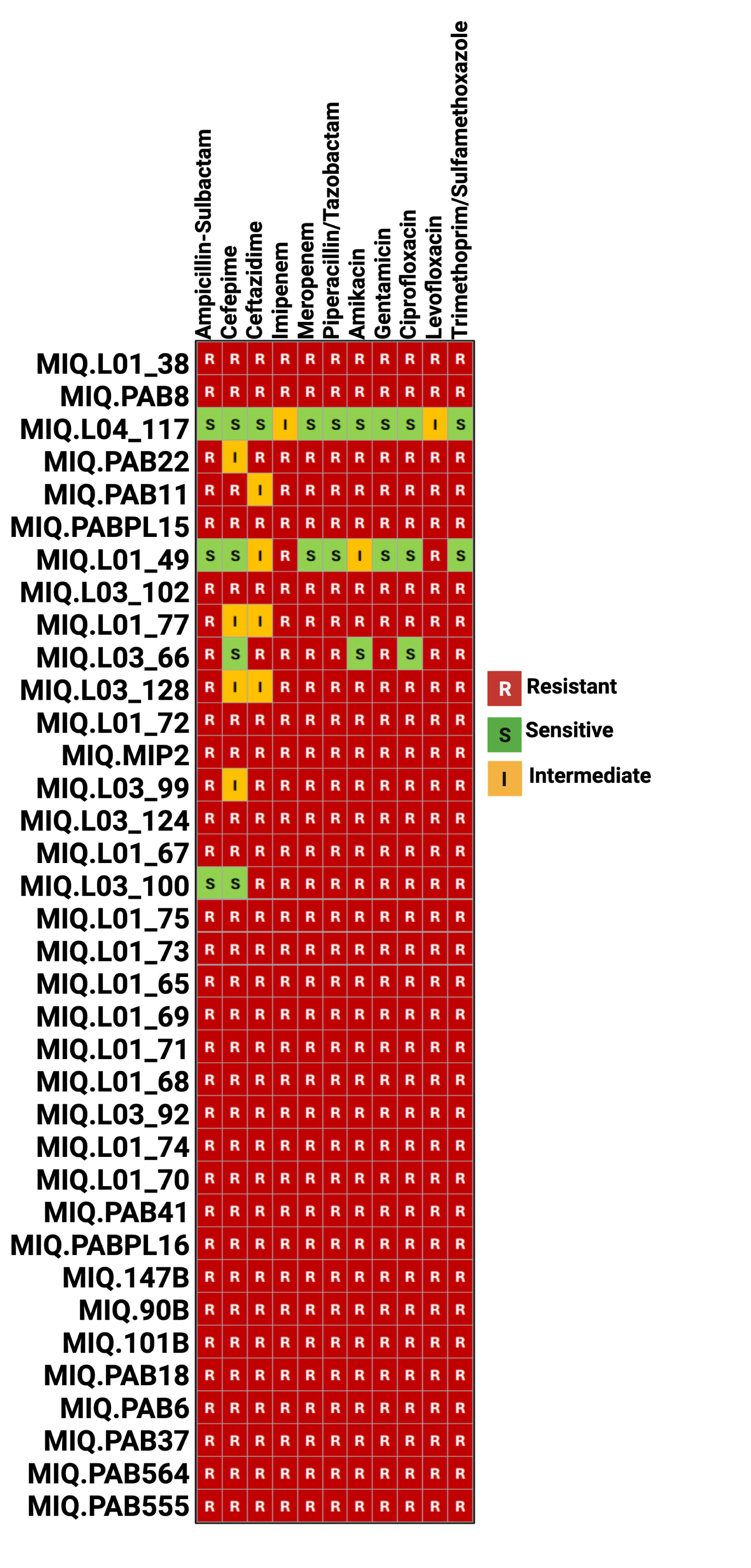
Phylogenetic landscape of *A. baumannii* in Mexico. (A) Maximum likelihood phylogenetic tree of 194 genomes, including 47 newly sequenced MIQ isolates and 147 historic published genomes (HPG). Clades A and B are subdivided into subclades (A.1–A.4, B.1–B.2), annotated with Oxford and Pasteur MLST sequence types. The outer ring indicates geographic origin (Jalisco, Aguascalientes, Mexico City, and other states). (B) Geographic distribution of isolates across Mexico, with counts per state. (C) Simplified tree showing the two dominant clades. Together, these panels illustrate lineage diversity, inter-regional mixing, and the persistence of epidemic clones across time.

Based on the canonical *gdhB* allele, the most frequent <u>Oxford STs</u> in our dataset (n=47 MIQ strains) were ST208 (24.0%), ST369 (11.3%), ST417 (19.1%), ST758 (26.0%), and ST136 (3.4%). Post-COVID isolates introduced rare and previously unreported sequence types, signaling new epidemiological trajectories (Figure 1). Within our MIQ collection, 6.3% could not be typed, and rare types such as STs 1321, 490, 227, 934, 1942, 690 and 225 appeared.

<u>Pasteur MLST</u> identified nine sequence types: ST2 (65%), ST156 (27%), ST1, ST25, ST79, ST193, ST203, ST318, ST422, and ST1544. ST2 (IC2) and ST156 (CC79, IC5) were the most prevalent, consistent with prior studies. ST2 was widely distributed across central-western Mexico (Aguascalientes, Jalisco, Mexico City, and sporadically Nuevo León, Coahuila, San Luis Potosí). ST156 grouped isolates from central and western Mexico, suggesting inter-regional dissemination.

Geographically, subclade A.1 corresponded to ST417 from Guadalajara and San Luis Potosí (Figure 1). Subclade A.2 was predominantly composed of ST369 isolates from Aguascalientes, Guadalajara, and Mexico City. Subclade A.3 was dominated by ST208 isolates from Guadalajara and Aguascalientes, with one isolate from Nuevo León. Subclade A.4 included ST136 (Guadalajara) and ST473 (Aguascalientes), with an additional isolate from Coahuila. Overall, Clade A demonstrated clear geographic mixing, prevalent between Jalisco and Aguascalientes.

Clade B was primarily composed of ST758 isolates (Figure 1). Subclade B.2 included isolates predominantly from Mexico City, whereas Subclade B.1 comprised isolates from Aguascalientes, Guadalajara, and Mexico City, suggesting regional dissemination of this lineage.

Three genomes (MIQ.90B, MIQ.PAB3, MIQ.PAB8) could not be assigned to existing Pasteur STs, suggesting novel emerging lineages. These isolates from Baja California and Guerrero, exhibited greater genetic heterogeneity. A closer comparative analysis showed that their closest related genomes were *Acinetobacter sp.* KPC-SM-2 (Germany, 2021), and *Acinetobacter baumannii* strain K09-14 (Malaysia, 2017).

### Resistome Profiles

Resistome profiling identified 128 distinct gene patterns and 44 emerging ARGs (Figure 2) in the HPG-MIQ dataset.

**Figure 2.**
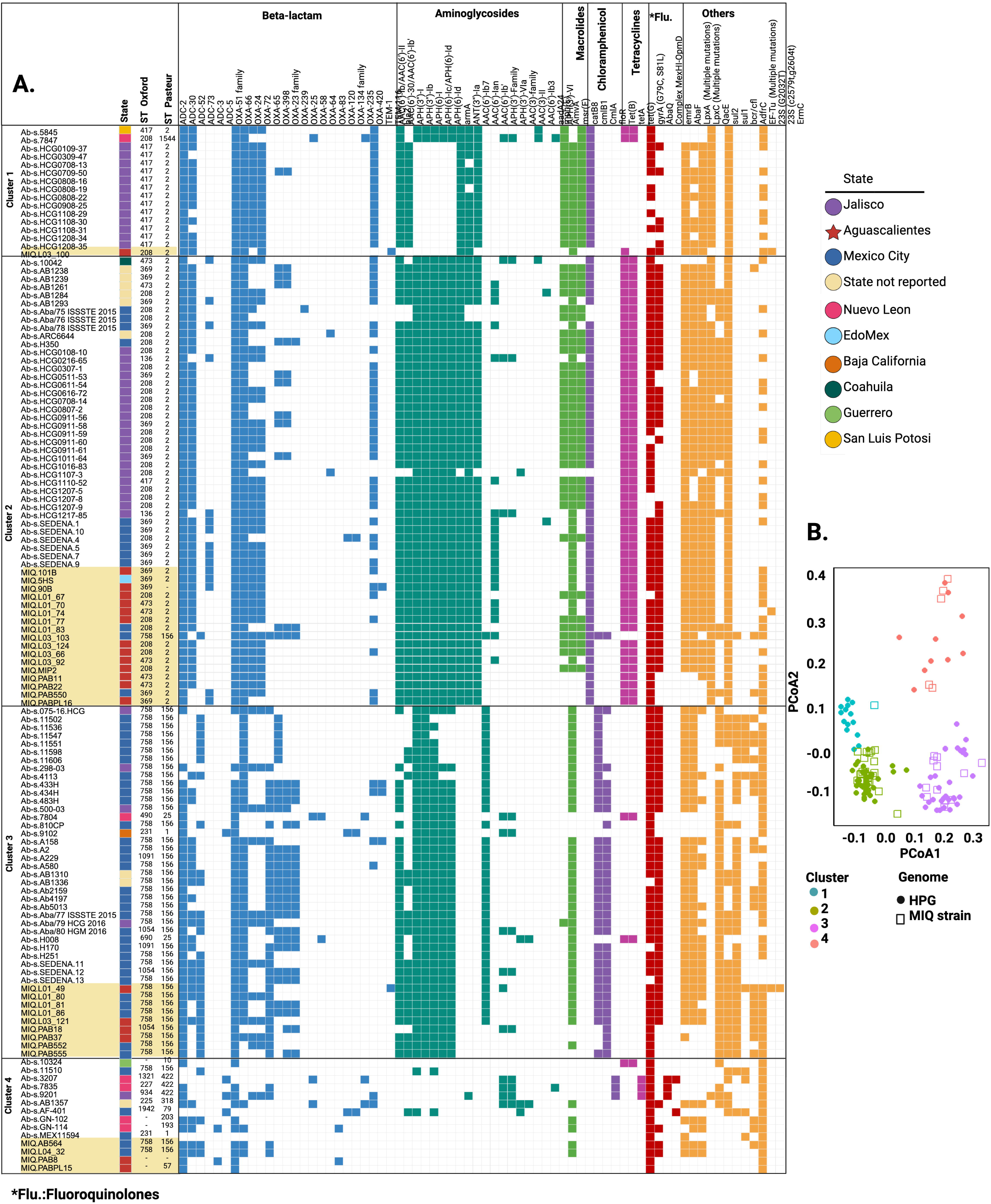
Resistome profiles. Heatmap showing antibiotic resistance gene (ARG) presence/absence across MIQ and HPG genomes. Genes are grouped by antibiotic class (β-lactams, aminoglycosides, macrolides, tetracyclines, fluoroquinolones, chloramphenicol, sulfonamides, trimethoprim, oxazolidinones, lipopeptides). Rows represent individual genomes, annotated with sequence type and state of origin. Columns represent ARGs; colored squares indicate presence. The diversity of resistome profiles highlights both conserved core genes and emerging ARGs enriched in post-COVID isolates.

The core resistome comprises 12 ARGs, including complex and target gene mutation. Genes with a unique presence in maximum two genomes (44 genes) (Table 2) were identified in five MIQ strains: MIQ.L01_49, MIQ.L03_100 AB54, MIQ.SA43a, MIQ.PAB3, and MIQ.PABPL15, as well as in some previously characterized strains.

**Table 1.**
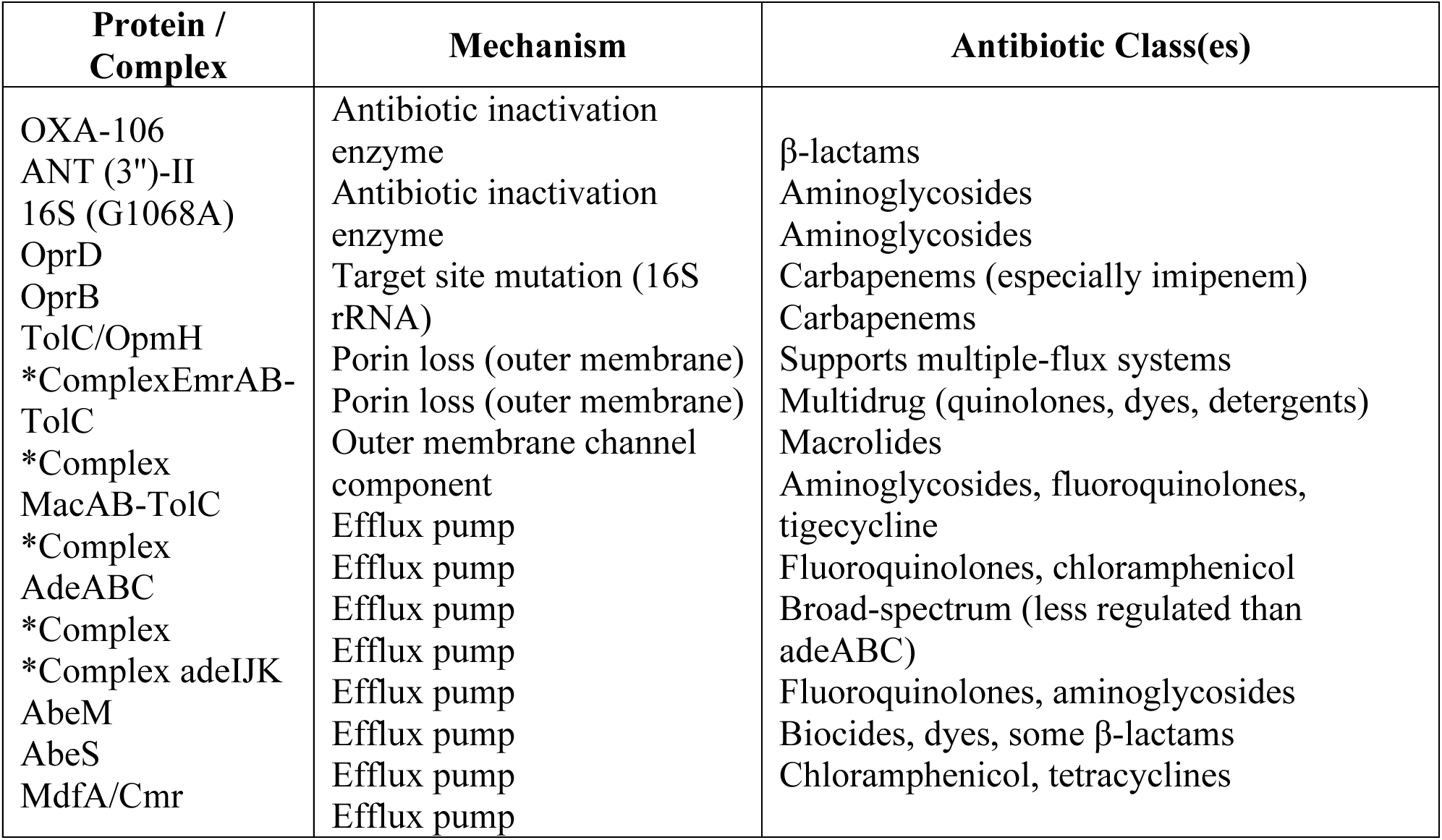
Antibiotic resistance genes (ARGs) identified in at least 188 out of 194 *A. baumannii* genomes analyzed (core ARGs). For gene families or efflux complexes, at least one representative gene was detected in each genome.

**Table 2.**
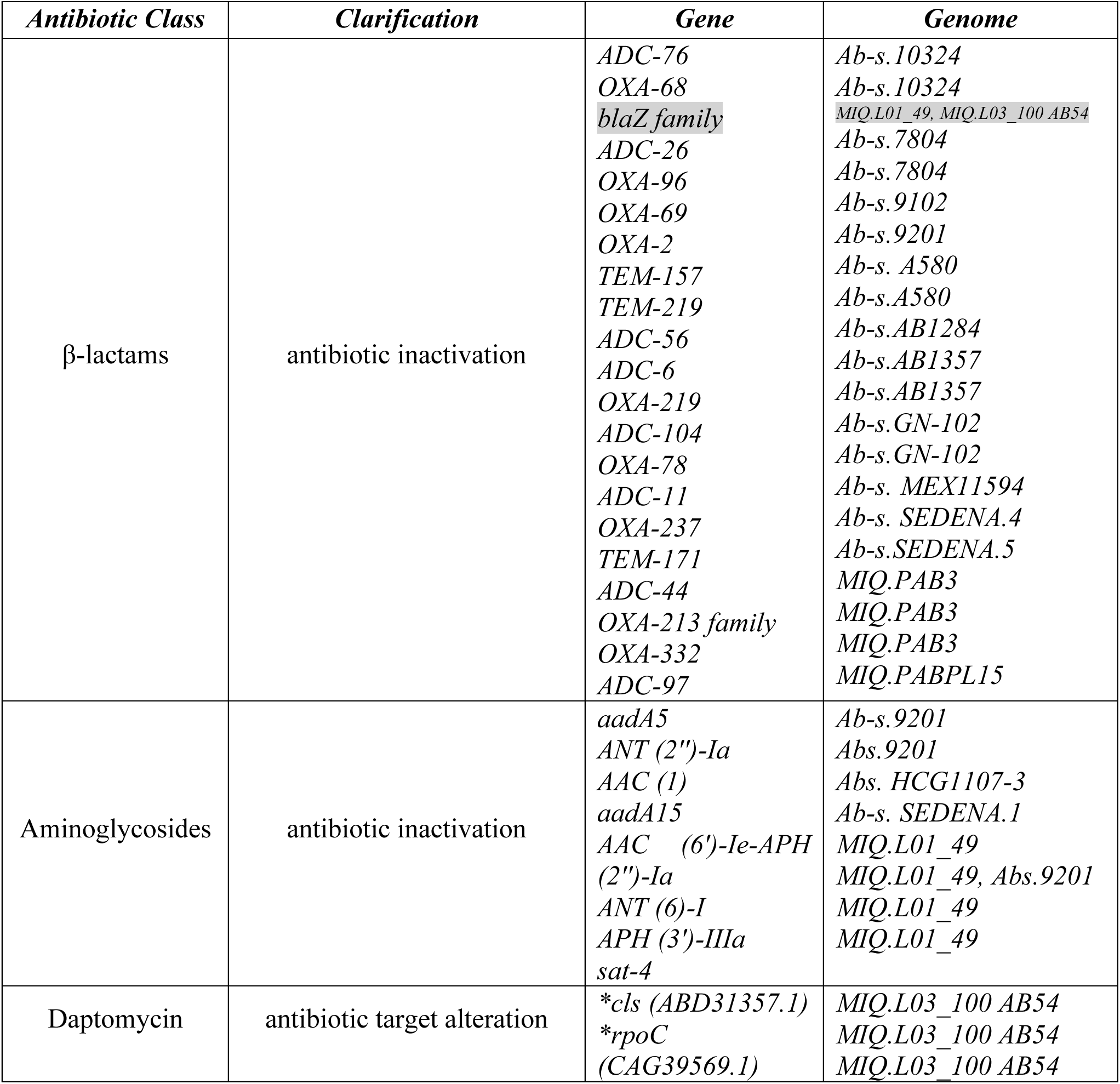
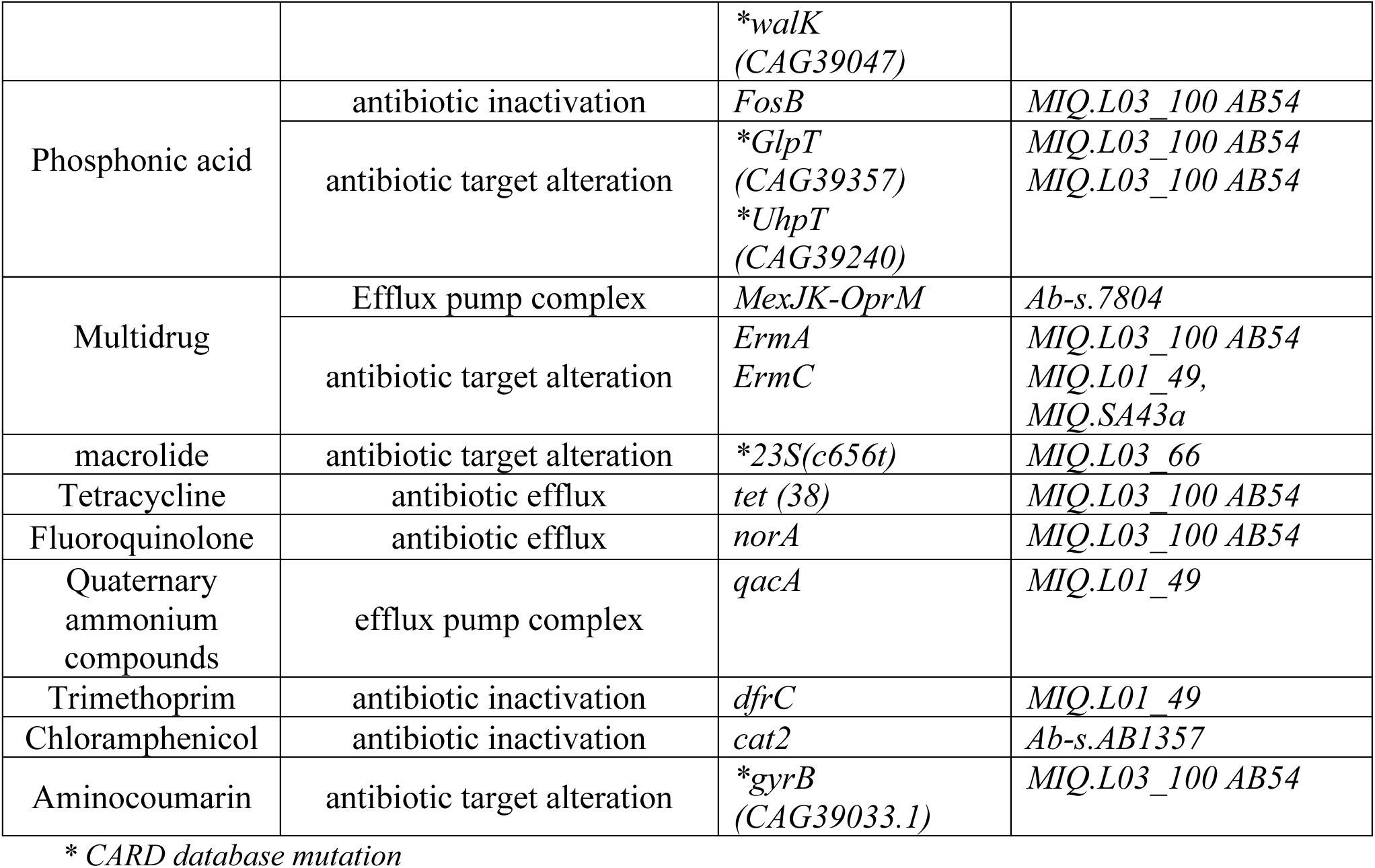
Antibiotic resistance genes (ARGs) identified in maximum 3 genomes.

### β-lactamases

A total of 50 β-lactamase genes were identified, including six ADC-type, twelve OXA-type, and four TEM-type variants not reported in the previous nationwide studies. Clusters enriched for carbapenemase genes (*blaOXA-72*, *blaOXA-66*) are overrepresented in referred MIQ isolates.

ADC-44 and ADC-97 were detected in MIQ.PAB3 and MIQ.PABPL15, respectively. Newly detected OXA-type β-lactamases, typically associated with *Acinetobacter calcoaceticus* (Figueiredo et al., 2012), included members of the OXA-213 family in MIQ.PAB3 and MIQ.L04_57, and OXA-332 exclusively in MIQ.PAB3. Other OXA and TEM variants were restricted to previously characterized genomes.

### Aminoglycosides

Resistance was mediated by 28 genes, including nine aminoglycoside acetyltransferases (*aac*) present in both strain groups. Three aminoglycoside adenyltransferases (*aad* genes) were exclusive to previously characterized strains. Four aminoglycoside nucleotidyltransferases (*ant*) were detected, with ANT(3’’)-II present in all strains.

Aminoglycoside phosphotransferases (*aph*) were distributed across clusters 1–3 but absent in cluster 4, which exhibited the greatest ST diversity. The *armA* gene, encoding an aminoglycoside methyltransferase, was restricted to clusters 1 and 2.

### Macrolides

Resistance genes *amvA, mphE*, and *msrE* were detected across clusters, while *ermA* and *ermC* were restricted to MIQ strains.

### Tetracyclines

Genes *tetB, tetA, tetG,* and *tet38* were identified exclusively in MIQ.L04_100 AB54.

### Chloramphenicol

Resistance determinants *catB8, cmlB1, cmlA,* and *floR* were detected across strains.

*Fluoroquinolones*.

Resistance was associated with mutations in *gyrA* (G79C, S81L) and the presence of the *adeFGH* efflux complex in all strains. Additional efflux determinants *abaQ, emrAB-tolC*, and *emrB* were exclusive to Ab-s.3207 and Ab-s.AF-401, while *norA* was unique to MIQ.L03_100 AB54.

### Sulfonamides and Trimethoprim

Genes *sul1* and *sul2* were widespread, whereas trimethoprim resistance mediated by *dfrC* was limited to clusters 3 and 4.

### Oxazolidinones and Lipopeptides

Linezolid resistance was linked to mutations in 23S rRNA (G2032T), with additional variants (C2579T, G2604T) detected in three strains, suggesting recent adaptive events. Daptomycin resistance-associated mutations in *walK* and *cls* were identified exclusively in MIQ.L03.100.

### Virulome Profiles

Virulence profiling revealed 19 distinct virulome patterns (Figure 3), with six exclusive to MIQ strains, eight exclusive to previously characterized strains, and five shared. Post-COVID isolates carried novel biofilm and outer-membrane protein genes, suggesting enhanced persistence in hospital environments and greater potential for immune evasion. Emerging virulence determinants were observed in strains MIQ.L03_100, MIQ.MIP2, MIQ.L01_71, MIQ.L01_49, and MIQ.AB550. Phylogenetic analysis showed no correlation between virulence gene distribution and phylogeny.

**Figure 3.**
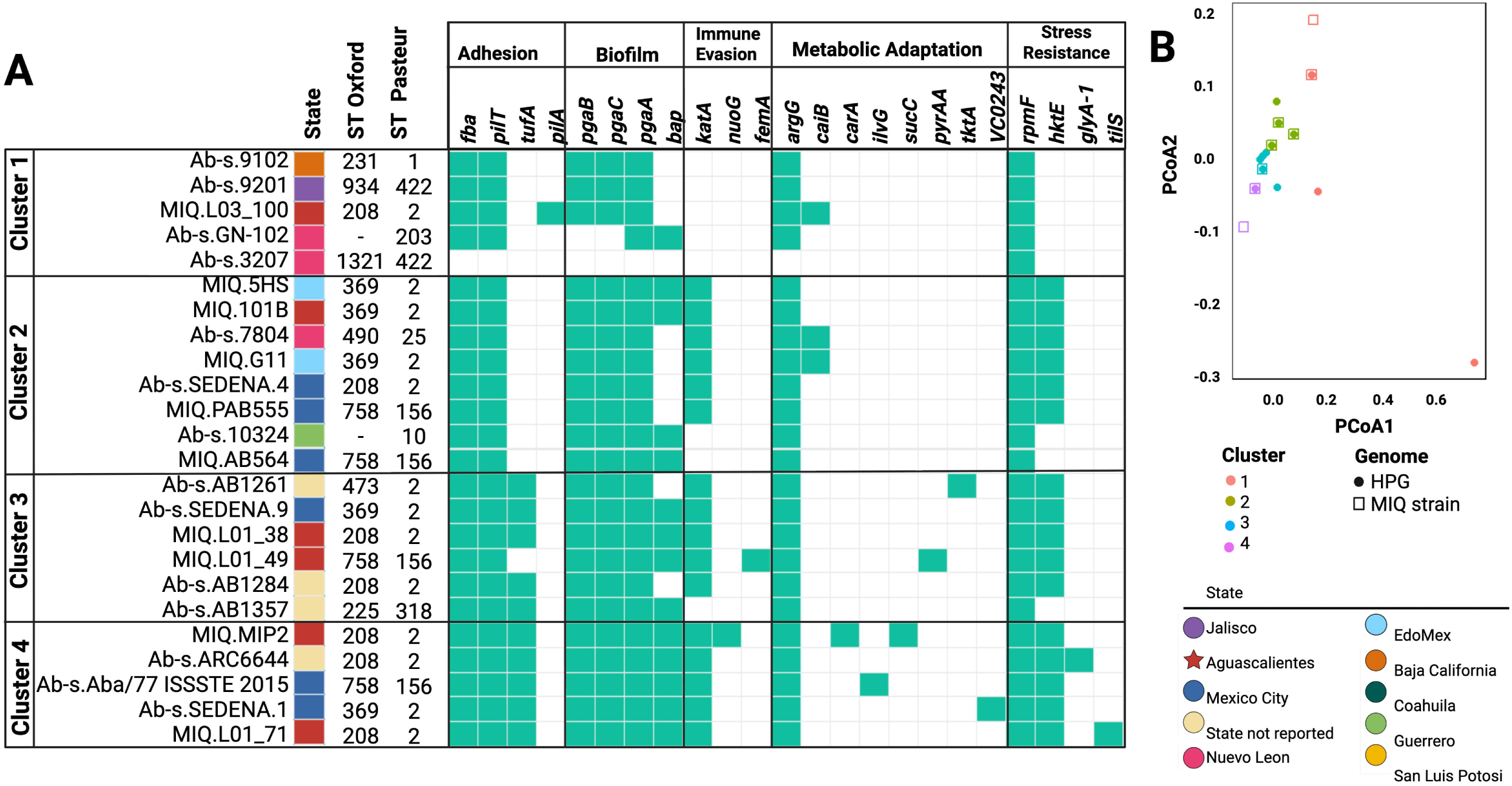
Virulome profiles. (A) Heatmap of virulence gene presence/absence across genomes, grouped into functional categories (adhesion, biofilm, immune evasion, metabolic adaptation, stress resistance). Rows represent strains, annotated with ST and state. Columns represent individual genes; green squares indicate presence. (B) Principal Coordinates Analysis (PCoA) based on Bray–Curtis dissimilarity of virulome profiles, showing four main clusters. Shapes distinguish MIQ and HPG genomes, colors correspond to clusters. Together, these panels demonstrate parallel evolution of virulence traits and their distribution across lineages.

We could identify four main virulome clusters, further supported by the PcoA analysis (Figure 4). Shared adhesion genes included *bap*, *pgaA*, *pgaB*, and *pgaC*, with notable copy number variation in *bap* (23 copies in Ab-s.H350 vs. 8 in MIQ.L01_38). Genes *pilT*, *fba*, and *tufA* were also common, while *pilA* was exclusive to MIQ.L03_100. Oxidative stress gene *katA* was present in both groups, whereas immune evasion genes *nouG* and *femA* were unique to MIQ strains. Metabolic genes *carA*, *sucC*, and *pyrAA* were exclusive to MIQ isolates, while *ilvG*, *glyA-1*, *tktA*, and *VC0243* were restricted to previously characterized strains. Additional findings included *hktE* and *rpmF* in both groups, *tilS* in MIQ.L01_71, and *glyA-1* in Ab-s.ARC6644.

**Figure 4.**
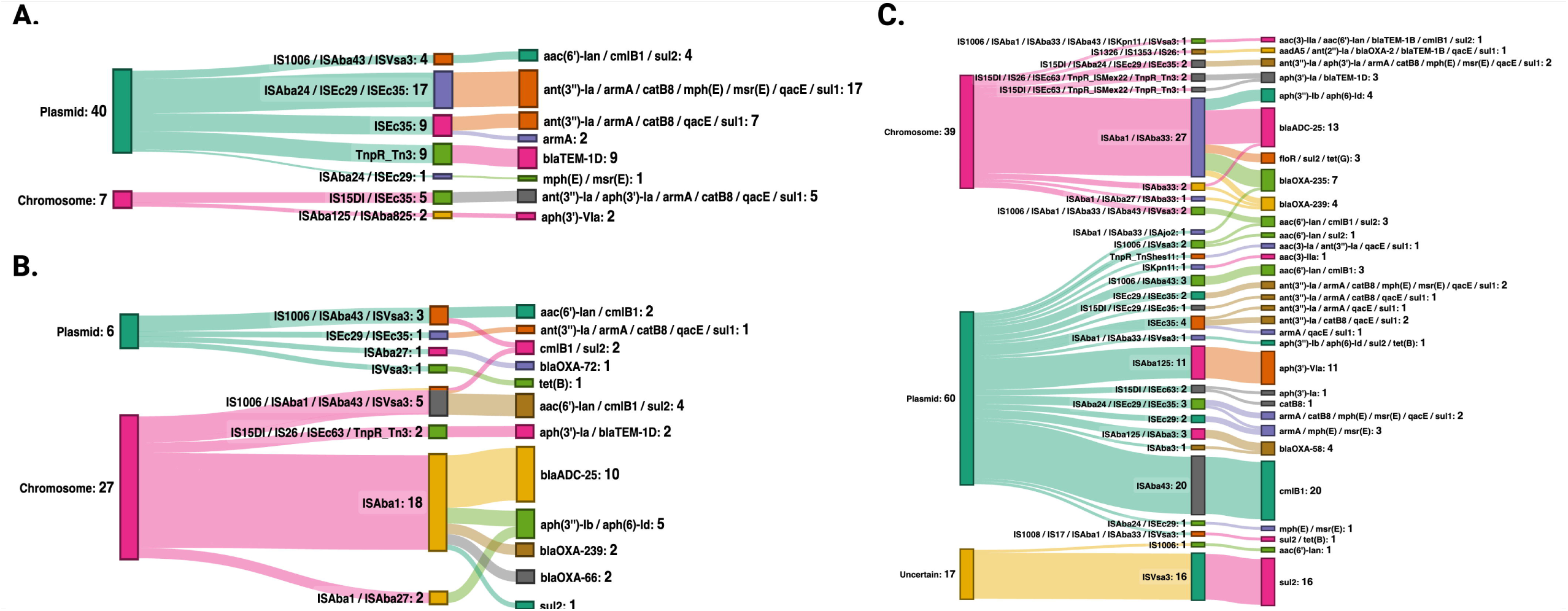
Mobilome composition. (A–C) Sankey diagrams illustrating gene flow between plasmids, chromosomes, and mobile genetic elements (MGEs) in MIQ and HPG genomes. Connections show associations between insertion sequences, transposons, and resistance genes. (D) Distribution of 24 MGEs identified across genomes. The diagrams highlight plasmids as the principal vectors of resistance dissemination and recurrent IS–ARG associations, including novel contexts in MIQ strains.

These results demonstrate that virulence traits evolved in parallel with resistance, generating pathogens that are both harder to treat and more persistent in clinical settings.

### Mobilome Composition

Plasmids were confirmed as the principal vectors of resistance gene dissemination, with 74.2% of MGEs plasmid-associated in MIQ strains, comparable to 76.7% in previously characterized isolates (Figure 5).

**Figure 5.**
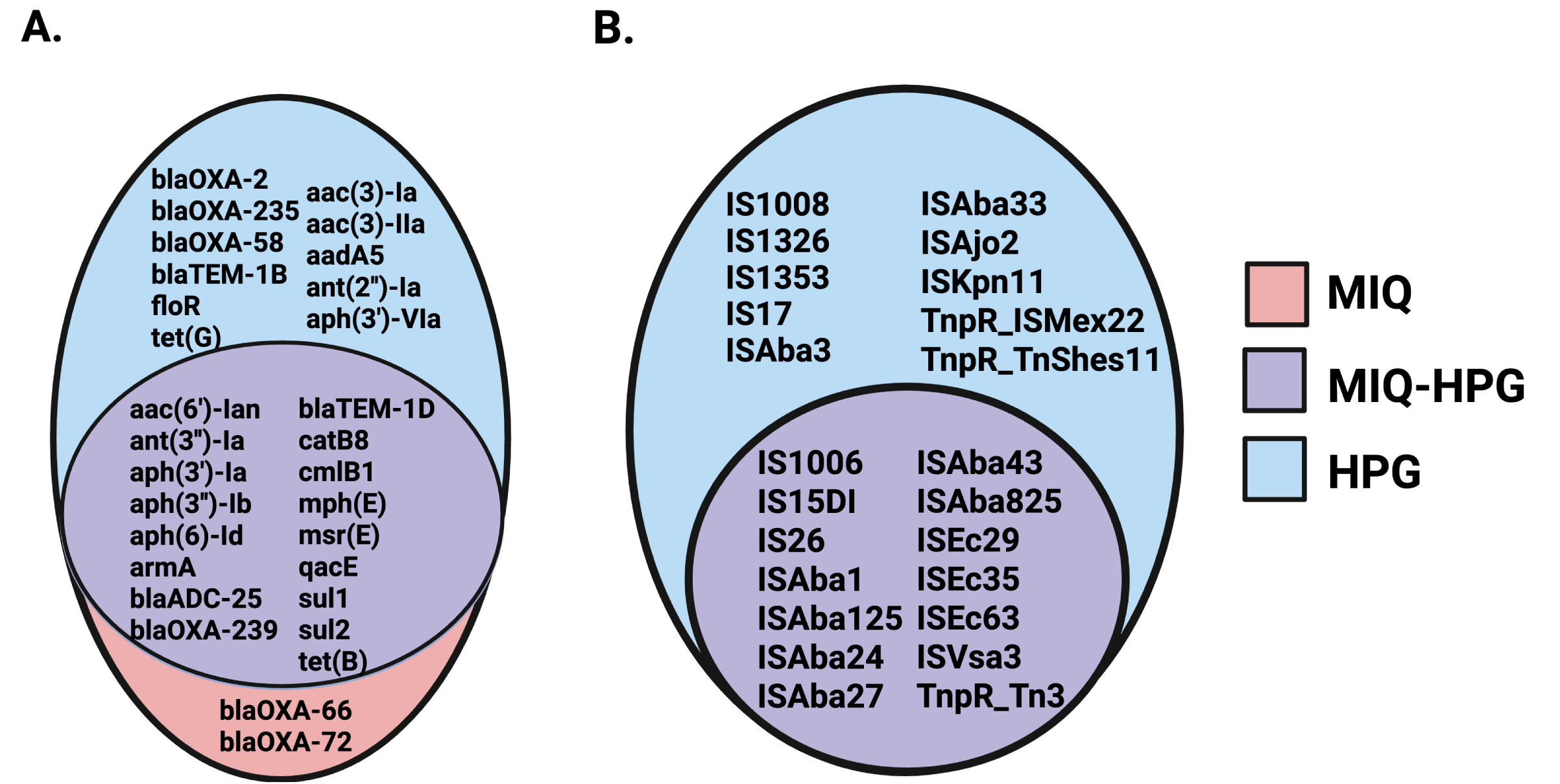
Comparative distribution of resistance genes and MGEs. (A) Venn diagram comparing resistance genes among MIQ, MIQ-HPG, and HPG genomes. Unique and shared ARGs are indicated in each set. (B) Venn diagram comparing insertion sequences and transposons among the same groups. These comparisons reveal conserved elements, lineage-specific repertoires, and novel associations in MIQ strains.

A total of 24 distinct mobile genetic elements (MGEs) were identified (Figure 4D) for the entire dataset. No novel MGEs were uniquely associated with MIQ strains; instead, these isolates revealed conserved MGEs linked to antimicrobial resistance gene (ARG) dissemination. In contrast, 30 different ARGs were found in genomic neighborhoods associated with MGEs, underscoring the diversity of resistance genomic context.

Recurrent structures included *IS1006*-associated modules carrying *sul2*, *cmlB1*, and *aac(6’)-Ian*, as well as multidrug resistance clusters involving *ISEc29* and *ISEc35* with *armA*, *ant(3’’)-Ia*, *catB8*, *mph(E)*, *msr(E)*, *qacE*, and *sul1*. Similarly, *ISAba1* and *ISAba33* were consistently associated with β-lactamase genes such as *blaADC-25* and *blaOXA-239*, in agreement with previously described genomic contexts.

Special attention was given to transposon-associated recombinases, particularly *TnpR* elements. These were identified in association with β-lactamase genes, including *TnpR_Tn3* linked to *blaTEM-1D* and *TnpR_TnShes11* linked to *aac(3)-Ia*, *ant(3’’)-Ia*, *qacE*, and *sul1*.

Notably, two β-lactamase genes were detected in novel insertion sequence contexts among MIQ isolates. The carbapenemase gene *blaOXA-72* was associated with *ISAba27*, while *blaOXA-66*—typically considered an intrinsic OXA-51-like variant—was identified in association with *ISAba1*. This represents a distinct genetic arrangement not observed in previously reported structures within this dataset. These findings highlight the emergence of new IS–ARG associations in MIQ strains, particularly involving clinically relevant OXA-type β-lactamases.

Comparative analysis of resistance genes and mobile genetic elements (MGEs) revealed distinct distributions between MIQ, MIQ-HPG, and HPG strains (Figure 6). In MIQ isolates, *blaOXA-66* and *blaOXA-72* were uniquely detected, while MIQ-HPG strains carried a broader repertoire including *aac(6’)-Ian*, *armA*, *blaADC-25*, *blaOXA-239*, *blaTEM-1D*, *catB8*, *cmlB1*, *mph(E)*, *msr(E)*, *qacE*, *sul1*, *sul2*, and *tet(B)*. HPG strains, in contrast, harbored *blaOXA-2*, *blaOXA-235*, *blaOXA-58*, *blaTEM-1B*, *floR*, *tet(G)*, and aminoglycoside resistance genes such as *aac(3)-Ia*, *aac(3)-IIa*, *aadA5*, *ant(2’’)-Ia*, and *aph(3’)-VIa*.

**Figure 8.**
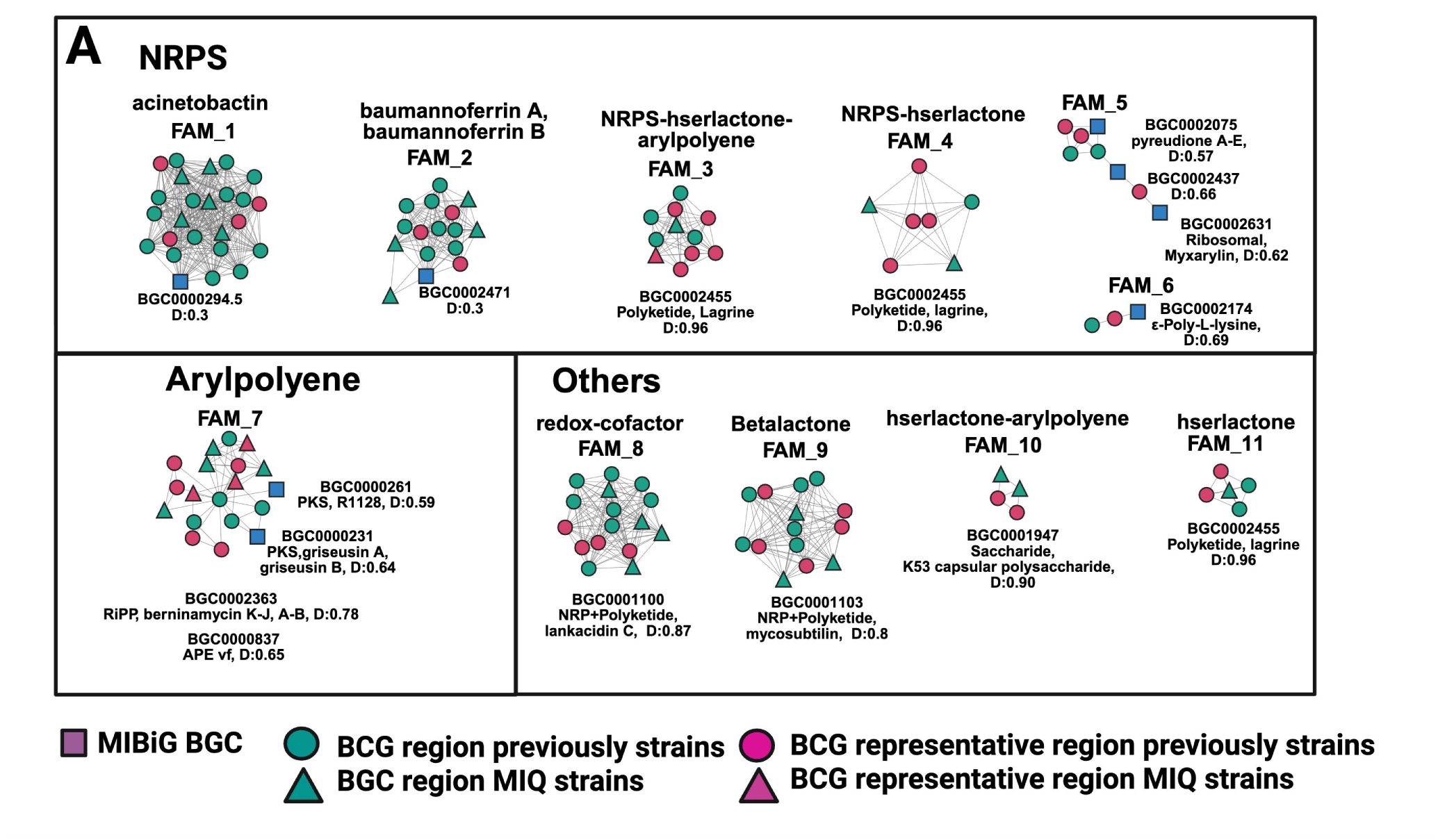
Biosynthetic gene clusters (BGCs) (A) Similarity network of 120 unique BGCs grouped into 11 families (FAM_1–FAM_11) using BiG-SCAPE. Nodes represent representative regions from MIQ and HPG genomes; edges indicate similarity (distance < 0.3). Families include conserved siderophores (acinetobactin, baumannoferrin A/B) and diverse NRPS, PKS, betalactone, and hserlactone clusters. The network highlights conserved iron acquisition systems and novel BGCs with antimicrobial potential in post-COVID isolates.

Insertion sequence profiling showed MIQ-HPG strains enriched for *IS1006*, *IS15DI*, *IS26*, *ISAba1*, *ISAba125*, *ISAba24*, *ISAba27*, *ISAba43*, *ISAba825*, *ISEc29*, *ISEc35*, *ISEc63*, *ISVsa3*, and *TnpR_Tn3*. In contrast, HPG strains carried *IS1008*, *IS1326*, *IS1353*, *IS17*, *ISAba3*, *ISAba33*, *ISAjo2*, *ISKpn11*, *TnpR_ISMex22*, and *TnpR_TnShes11*. These distributions highlight both conserved and lineage-specific mobilome structures, with MIQ strains showing novel associations involving clinically relevant OXA-type β-lactamases.

### Specialized Metabolism

Secondary metabolite analysis revealed conserved siderophore biosynthetic clusters across all clades, underscoring iron acquisition as a central survival strategy. Post-COVID isolates also carried novel biosynthetic gene clusters (BGCs) with antimicrobial potential, suggesting unexplored metabolic capacities that may influence interspecies competition and host interactions.

Using antiSMASH 7.0, we identified 1,637 genomic regions linked to specialized metabolite production across 204 genomes, with an average of eight BGCs per genome; strain Ab-s.SEDENA6 carried 11. Clustering with BiG-SCAPE 2.0 reduced redundancy, selecting 42 representative regions and yielding 120 unique BGCs grouped into 11 families (distance < 0.3) (Figure 7).

Six families (FAM_1–FAM_6) encoded nonribosomal peptide synthetases (NRPS), including *baumannoferrin A/B* and *acinetobactin* (FAM_1, FAM_2), present in most strains. FAM_3 and FAM_4 represented hybrid NRPS-hserlactone-arylpolyene clusters, while FAM_5 and FAM_6 showed high genomic variation. FAM_7 produced arylpolyene PKS clusters related to *berninamycins* and *APE Vf*. FAM_8 encoded a redox cofactor cluster similar to *lankacidin C*, FAM_9 a betalactone cluster related to *mycosubtilin*, FAM_10 a hserlactone-arylpolyene cluster resembling the K53 capsular polysaccharide locus, and FAM_11 a hserlactone producer.

## Discussion

Our integrative genomic analysis demonstrates that the COVID-19 pandemic reshaped the genomic architecture of multidrug-resistant *Acinetobacter baumannii* in Mexico. By analyzing 194 genomes, including 47 newly sequenced post-pandemic isolates (MIQ), we identified two major clades dominated by Oxford STs 758, 208, 417, and 369. These findings align with prior national studies reporting distinctive sequence type distributions in Mexico compared to neighboring countries (Barrios-Camacho et al., 2025; Fernández-Vázquez et al., 2023). The persistence of Pasteur ST2 and ST156 confirms the dominance of international clones IC2 and IC5, which are widely associated with carbapenem resistance and high virulence (Gaiarsa et al., 2019; Shelenkov et al., 2023; Boutzoukas and Doi, 2025). IC2 is the most widespread lineage globally, while IC5 is particularly enriched in Latin America, including Mexico, Brazil, and Paraguay (Pearl and Anbarasu, 2024). Other epidemic clones such as IC1, IC7, and IC8 dominate in Asia and Europe but remain rare in Mexico (Sholeh et al., 2025).

The emergence of rare and previously unreported Oxford STs (e.g., ST1321, ST490, ST227, ST934, ST1942, ST690, ST225) suggests ongoing diversification and potential introduction of novel lineages. These patterns likely reflect inter-hospital or inter-state transmission routes facilitated by patient transfers or shared healthcare practices, underscoring the need for coordinated surveillance across regions rather than isolated hospital-level monitoring (Barrios-Camacho et al., 2025).

Resistome profiling revealed 128 distinct gene patterns and 44 emerging ARGs, expanding the repertoire beyond classical carbapenemases and efflux pumps. Their presence suggests interspecies gene transfer facilitated by mobile genetic elements (Fernández-Vázquez et al., 2023; Wick et al., 2023). Carbapenemase gene enrichment (*blaOXA-72*, *blaOXA-66*) in MIQ strains was likely influenced by referral bias, as hospital laboratories preferentially submit resistant isolates. Policymakers should therefore interpret hospital-based genomic surveillance cautiously and complement it with community-level sampling (Manobanda Nata and Jaramillo Ruales, 2023).

Virulome analysis confirmed conserved adhesion, biofilm, and immune evasion genes, consistent with previous reports (Sholeh et al., 2023). However, MIQ strains exhibited expanded repertoires of metabolic adaptation genes, suggesting enhanced survival under stress conditions. This adaptation may contribute to persistence in hospital environments, where antibiotic pressure and host immune responses create challenging niches.

Mobilome analysis highlighted plasmid-mediated dissemination of ARGs, with enrichment of *ISAba1* and *ISAba3* elements in post-COVID isolates. These insertion sequences are known to activate carbapenemase genes and facilitate horizontal gene transfer (Wick et al., 2023). Even though assemblies were fragmented, the presence of these MGEs indicates ongoing plasmid-mediated ARG dissemination. Their increased prevalence in recent isolates underscores the urgency of plasmid-focused surveillance using long-read sequencing (Fernández-Vázquez et al., 2023). Novel IS–ARG contexts, such as *ISAba27–blaOXA-72* and *ISAba1–blaOXA-66*, highlight local innovation in mobilome dynamics, paralleling global reports of diverse IS–ARG associations (Chukamnerd et al., 2025; Ristori et al., 2025).

Specialized metabolism analysis revealed conserved siderophores (*acinetobactin*, *baumannoferrin*) alongside novel biosynthetic clusters (Djahanschiri et al., 2022; Djahanschiri et al., 2024). Expanded BGC repertoires may enhance interspecies competition and persistence in polymicrobial infections, complicating treatment outcomes. This study represents the first genomic survey of specialized metabolism in Mexican *A. baumannii*, highlighting both conserved siderophore systems and novel BGCs with potential roles in virulence, biofilm stability, and antimicrobial activity.

Together, these findings provide the first integrative genomic portrait of *A. baumannii* in Mexico across the pre- and post-COVID periods. They situate Mexican isolates within the global CRAb crisis, dominated by IC2 and IC5, but also reveal unique mobilome innovations and novel IS–ARG contexts. Sustained genomic surveillance, antimicrobial stewardship, and functional validation of genomic predictions are urgently needed. Without these measures, continued diversification of *A. baumannii* lineages and expansion of resistance determinants may undermine infection control efforts and therapeutic efficacy.

## Conclusion

This study provides the first integrative genomic portrait of *Acinetobacter baumannii* circulating in Mexico across the pre- and post-COVID-19 periods. By analyzing 194 genomes, including 47 newly sequenced isolates, we revealed how resistance determinants, virulence factors, mobile genetic elements, and biosynthetic gene clusters converged to shape novel pathogenic lineages. The dominance of Oxford STs 758, 208, 417, and 369 highlights the persistence of internationally recognized clones, while the emergence of rare sequence types signals ongoing local diversification.

Beyond these core findings, several additional observations carry important public health relevance. Phylogenetic analyses demonstrated geographic mixing of clades across Jalisco, Aguascalientes, and Mexico City, suggesting inter-hospital or inter-state transmission routes that demand coordinated regional surveillance rather than isolated hospital-level monitoring.

Resistome profiling revealed referral bias toward carbapenemase-positive isolates, underscoring the need to complement hospital-based genomic surveillance with community-level sampling to avoid overestimating resistance prevalence. Virulome analysis showed enrichment of metabolic adaptation genes in MIQ strains, pointing to enhanced resilience that may facilitate persistence in hospital environments and increase outbreak risk. Mobilome profiling identified enrichment of ISAbA1 and ISAbA3 elements in post-COVID isolates, highlighting ongoing plasmid-mediated dissemination of carbapenemase genes and the urgency of incorporating long-read sequencing into surveillance frameworks. Finally, biosynthetic gene cluster analyses revealed expanded repertoires in MIQ genomes, suggesting adaptive metabolic diversification that may enhance ecological fitness and complicate treatment outcomes.

Taken together, these findings emphasize that *A. baumannii* in Mexico is not only evolving through classical resistance mechanisms but also through geographic mixing, referral biases, virulome expansion, mobilome diversity, and specialized metabolism. Addressing these challenges requires integrated genomic surveillance, functional validation of adaptation signals, and regionally coordinated infection control strategies. Without such measures, the continued diversification of *A. baumannii* lineages and expansion of resistance determinants may undermine therapeutic efficacy and infection control efforts in the post-COVID era.

## Author Contributions

Corina-Diana Ceapă: Conceptualization, Supervision, Writing – original draft, Writing – review & editing Moisés A. Alejo: Data curation, Investigation, Visualization Karla G. Hernández Magro Gil: Methodology, Validation Miriam Sarahi Lozano Gamboa: Clinical sampling, Logistics Uriel Colin Camacho: Bioinformatics, Data analysis Brian Muñoz Gomez: Laboratory work, Sample processing Rodolfo García-Contreras: Review, Interpretation Rachel J. Whitaker: Comparative genomics, Review Anastasio Palacios Marmolejo: Metadata curation, Ethics compliance

## Funding

This work was supported by collaborations with MGI and Illumina for strain sequencing at no cost to the research group. We acknowledge receiving funding from the Illinois Mexican & Mexican American Students Initiative (I-MMAS) and the Coordinación de la Investigación Científica of UNAM Seed Funding 2024 for the project “Targeting of mobile elements from multidrug-resistant ESKAPEE pathogens”. The funders had no role in study design, data collection and analysis, decision to publish, or preparation of the manuscript.

## Acknowledgments

We thank the staff of the State Public Health Laboratory of Aguascalientes (LESPA) for their assistance in sample processing and logistics, and obtaining the ethican and biosafety permits. We acknowledge the technical support provided by the sequencing facilities at Roy J. Carver Biotechnology Center, University of Illinois at Urbana-Champaign. We would like to thank Applikon and Illumina, and in particular Ileana Gutierrez, Alejandra García, Verónica Sánchez, and Ethel Salazar, for including the laboratory in the 2023 sequencing course and training, which allowed us to sequence 94 bacterial genomes, data that were used to complete this work.

We would like to thank MGI and Química Valaner, in particular Blanca Mondragon, for their help in sequencing 92 strains of bacterial pathogens in 2022.

The figures were created in https://BioRender.com.

## Conflict of Interest

The authors declare that the research was conducted in the absence of any commercial or financial relationships that could be construed as a potential conflict of interest.

## Data Availability Statement

All genome assemblies generated in this study have been deposited in NCBI under BioProject accession number **PRJNA1291376**. Associated metadata, including collection sites, dates, and sources, are provided in Supplementary Table 1.

## Ethics Statement

This study was approved by the Promotora Médica Aguascalientes S.A. de C.V. Ethics and Biosafety Committee (Approval Nos. 100.cbpma.2025 and 2937.ceipma.2025), registered with COFEPRIS. Informed consent was obtained in accordance with Mexican regulations (NOM-012-SSA3-2012).

## Supplementary Material

Supplementary Table 1. Metadata of *Acinetobacter baumannii* isolates collected from hospitals, environmental sources, and reference strains across Mexico. The table includes GenBank accession numbers, genome identifiers, collection year, hospital/state of origin, sequence type (ST) assignments (Pasteur and Oxford schemes), patient demographics, comorbidity status, COVID-19 test/vaccine information, and clinical outcomes.

Supplementary Table 2. Distribution of antimicrobial resistance (AMR) gene families across *A. baumannii* isolates. Gene presence/absence is reported per genome, grouped by functional categories (aminoglycoside, β-lactam, macrolide, sulfonamide, tetracycline, etc.), enabling comparative analysis of resistance determinants between MIQ and historic strains.

Supplementary Table 3. Virulence gene distribution across *A. baumannii* isolates. The table details the presence of genes associated with adhesion, biofilm formation, iron acquisition, secretion systems, and other virulence traits, highlighting variability across clinical and environmental strains.

Supplementary Table 4. Mobile genetic elements (MGEs) identified in *A. baumannii* genomes. The table lists ARG-associated MGEs, including plasmids, integrons, transposons, insertion sequences (IS), genomic islands (GI), integrative conjugative elements (ICE), and prophages. Coordinates and replicon details are provided to illustrate the genomic context of resistance and virulence determinants.

Supplementary Table 5. Biosynthetic gene clusters (BGCs) identified in MIQ and historic *A. baumannii* strains. The table reports cluster types (e.g., siderophores, non-ribosomal peptide synthetases, polyketides), genomic coordinates, and predicted functions, enabling comparative analysis of secondary metabolite potential across isolates.

## Supplementary Methods

Detailed bioinformatics pipeline for resistome, virulome, mobilome, and BGC analyses.

## Notes

### Competing Interest Statement

The authors have declared no competing interest.

## References

Barrios-Camacho, B., Lozano-Aguirre, L., and Durán-Bedolla, J. (2025). Genomic analysis of the main epidemiological lineages of *Acinetobacter baumannii* in Mexico. Front. Cell. Infect. Microbiol. 14, 1499839. doi: 10.3389/fcimb.2024.1499839

Boutzoukas, A., and Doi, Y. (2025). Global epidemiology of carbapenem-resistant *Acinetobacter baumannii*. JAC-Antimicrob. Resist. 7:dlf123. doi:10.1093/jacamr/dlf123

Castillo-Bejarano, J.I., Llaca-Díaz, J., Mascareñas de los Santos, A.H., Sánchez-Alanís, H., Espinosa-Villaseñor, F., Aguayo-Samaniego, R., et al. (2023). Carbapenem-resistant *Acinetobacter baumannii* infection in children from a third-level hospital in Mexico: clinical characteristics and molecular epidemiology. J. Pediatric Infect. Dis. Soc. 12, 431–435. doi: 10.1093/jpids/piad046

Chapartegui-González, I., Lázaro-Díez, M., Redondo-Salvo, S., Navas, J., Ramos-Vivas, J. (2021). Antimicrobial resistance determinants in genomes and plasmids from *Acinetobacter baumannii* clinical isolates. Antibiotics 10(7). doi: 10.3390/antibiotics10070753

Chukamnerd, A., Wongchai, T., Srisamang, P., and Chantratita, N. (2025). Genomic epidemiology of carbapenem-resistant *Acinetobacter baumannii* in Thailand. Antibiotics 14:112. doi:10.3390/antibiotics14010112

Djahanschiri, B., Di Venanzio, G., Distel, J.S., Breisch, J., Dieckmann, M.A., Goesmann, A., et al. (2022). Evolutionarily stable gene clusters shed light on pathogenicity in the *Acinetobacter calcoaceticus-baumannii* complex. PLOS Genet. 18, e1010020. doi: 10.1371/journal.pgen.1010020

Djahanschiri, B., Di Venanzio, G., Breisch, J., Goesmann, A., Müller, J., et al. (2024). Biosynthetic gene cluster diversity in *Acinetobacter baumannii* and its role in adaptation. Front. Microbiol. 15, 1183927. doi: 10.3389/fmicb.2024.1183927

Fernández-Vázquez, J.L., Hernández-González, I.L., Castillo-Ramírez, S., Jarillo-Quijada, M.D., Gayosso-Vázquez, C., Mateo-Estrada, V.E., et al. (2023). Pandrug-resistant *Acinetobacter baumannii* from different clones and regions in Mexico share a plasmid carrying the blaOXA-72 gene. Front. Cell. Infect. Microbiol. 13, 1278819. doi: 10.3389/fcimb.2023.1278819

Figueiredo, S., Bonnin, R. A., Poirel, L., Duranteau, J., & Nordmann, P. (2012). Identification of the naturally occurring genes encoding carbapenem-hydrolysing oxacillinases from *Acinetobacter haemolyticus, Acinetobacter johnsonii*, and *Acinetobacter calcoaceticus*. Clinical microbiology and infection : the official publication of the European Society of Clinical Microbiology and Infectious Diseases, 18(9), 907–913. 10.1111/j.1469-0691.2011.03708.x

Gaiarsa, S., Batisti Biffignandi, G., Esposito, E.P., Castelli, M., Jolley, K.A., Brisse, S., et al. (2019). Comparative analysis of the two *Acinetobacter baumannii* MLST schemes. Front. Microbiol. 10, 930. doi: 10.3389/fmicb.2019.00930

Gomes Chagas, T.P., Rangel, K., and De-Simone, S.G. (2024). Carbapenem-resistant *Acinetobacter baumannii* in Latin America. In: Rangel, K., and De-Simone, S.G., eds. Acinetobacter baumannii – The Rise of a Resistant Pathogen. London: IntechOpen. doi: 10.5772/intechopen.1003713

Guillén-Navarro, D., Ochoa, S.A., De la Rosa-Zamboni, L., Giono-Cerezo, S., Xicohtencatl-Cortes, J., et al. (2025). Comparative genomics of carbapenem-resistant *Acinetobacter baumannii* isolated from pediatric patients in a tertiary care hospital. Microbiol. Spectr. 13(11). doi: 10.1128/spectrum.01676-25

Hernández-González, I.L., Mateo-Estrada, V., Castillo-Ramírez, S. (2022). The promiscuous and highly mobile resistome of *Acinetobacter baumannii*. Microb. Genom. 8(1). doi: 10.1099/mgen.0.000762

Huang L-Y, Lu P-L, Chen T-L, Chang F-Y, Fung C-P, Siu LK. (2010) Molecular Characterization of β-Lactamase Genes and Their Genetic Structures in *Acinetobacter* Genospecies 3 Isolates in Taiwan. Antimicrob Agents Chemother 54:2699–2703. doi: 10.1128/AAC.01624-09

Humberto, B.C., Luis, L.A., Josefina, D.B. (2025). Genomic analysis of the main epidemiological lineages of *Acinetobacter baumannii* in Mexico. Front. Cell. Infect. Microbiol. 14. doi: 10.3389/fcimb.2024.1499839

López-Jácome, L.E., Fernández-Rodríguez, D., Franco-Cendejas, R., Camacho-Ortiz, A., Morfin-Otero, M.D.R., Rodríguez-Noriega, E., et al. (2022). Increment antimicrobial resistance during the COVID-19 pandemic: results from the INVIFAR Network. Microb. Drug Resist. 28(3), 338–345. doi: 10.1089/mdr.2021.0231

Manobanda Nata, C.I., and Jaramillo Ruales, E.K. (2023). Carbapenem-resistant *Acinetobacter baumannii* complex: a review in Latin America. Salud, Ciencia y Tecnología 3, 479. doi: 10.56294/saludcyt2023479

Pan American Health Organization (PAHO/WHO). (2023). Antimicrobial Resistance (AMR) Portal: Surveillance of antimicrobial resistance in Latin America. Washington, DC: PAHO.

Panunzi, L.G. (2020). Corrigendum: sraX: A Novel Comprehensive Resistome Analysis Tool. Front. Microbiol. 11, 52. doi: 10.3389/fmicb.2020.594635

Pearl, S., and Anbarasu, A. (2024). Genomic landscape of nosocomial *Acinetobacter baumannii*: resistome, virulome, mobilome. Sci. Rep. 14:5678. doi:10.1038/s41598-024-56789

Ristori, M. V., Rossi, F., Piras, C., and Brisse, S. (2025). Genomic surveillance of *Acinetobacter baumannii* in Europe: diversity and dissemination of carbapenem-resistant clones. Microorganisms 13:456. doi:10.3390/microorganisms13030456

Rodrigues, D.L.N., Morais-Rodrigues, F., Hurtado, R., Dos Santos, R.G., Costa, D.C., Barh, D., et al. (2021). Pan-resistome insights into the multidrug resistance of *Acinetobacter baumannii*. Antibiotics 10(5). doi: 10.3390/antibiotics10050596

Shelenkov, A., Petrova, N., Mikhaylova, Y., and Akimkin, V. (2023). International clones of high risk *Acinetobacter baumannii*: definitions, history, properties and perspectives. Microorganisms 11:1234. doi:10.3390/microorganisms11061234

Sholeh, M., Rahimi, F., Azimi, L., and Pourmand, M. R. (2023). Virulence determinants and biofilm formation in multidrug-resistant *Acinetobacter baumannii*: global perspectives. Front. Microbiol. 14:765432. doi:10.3389/fmicb.2023.765432

Sholeh, M., Hamidieh, F., Beig, M., Badmasti, F. (2025). Unravelling the genomic landscape of *Acinetobacter baumannii*: deep dive into virulence factors, resistance elements, and evolutionary adaptations. Mol. Genet. Genomics 300(1), 68. doi: 10.1007/s00438-025-02265-3

Wang, M., Goh, Y.X., Tai, C., Wang, H., Deng, Z., Ou, H.Y. (2022). VRprofile2: Detection of antibiotic resistance-associated mobilome in bacterial pathogens. Nucleic Acids Res. 50(W1), W768–W773. doi: 10.1093/nar/gkac321

Wick, R.R., Judd, L.M., and Holt, K.E. (2023). Deep sequencing of *Acinetobacter baumannii* reveals complex mobilome architecture. Microb. Genom. 9, 000934. doi: 10.1099/mgen.0.000934

