## Supplementary Materials for "Resistome, Virulome, Mobilome, And Biosynthetic Gene Clusters Adaptations of *Acinetobacter Baumannii* Mexican Strains Across the Pre- and of the COVID-19 Period: Insights from Whole-Genome Sequencing"

### Supplementary Material: Extended Bioinformatics Analyses

#### Whole Genome Sequencing (WGS)

Genomic DNA from 47 newly sequenced A. baumannii isolates (2020–2023, post-COVID period) was extracted using the NEB Monarch® Genomic DNA Purification Kit with minor modifications. DNA integrity was confirmed by agarose gel electrophoresis, and purity assessed by NanoDrop™ spectrophotometry. Libraries were prepared with Illumina MiSeq™ DNA Prep v3, quantified with Qubit 1X dsDNA HS, pooled, denatured, and sequenced on an Illumina MiniSeq (paired-end 2×150 bp).

#### Genome Assembly and Annotation

Raw reads were quality-checked with FastQC v0.11.9 (Andrews, 2010), trimmed with Trimmomatic v0.39 (Bolger et al., 2014), and assembled *de novo* using SPAdes v3.9.0 (Bankevich et al., 2012). Assemblies were annotated with BV-BRC v3.30.19 (RAST toolkit).

#### Phylogenetic Analyses

Genomes were annotated with Prokka v1.14.6 (Seemann, 2014), core genome alignments generated with Roary v3.13.0 (Page et al., 2015), and maximum likelihood trees built with IQ-TREE v2.1.2 (Minh et al., 2020). Trees were visualized in iTOL v6 (Letunic and Bork, 2021).

#### Resistome and Virulome Profiling

Resistome annotation combined BV-BRC K-mer pipeline, CARD v3.2.9 (Alcock et al., 2020), NDARO (NCBI, 2023), and SraX (Panunzi, 2020). Virulome profiling integrated BV-BRC annotation with the VICTORS database (Sayers et al., 2019). Diversity was quantified using Bray–Curtis indices, visualized by PCoA, and validated with PERMANOVA.

**Mobilome Analysis**

Mobile genetic elements (MGEs) were identified using VRprofile2 (Wang et al., 2022). Sankey diagrams were generated with SankeyMATIC (SankeyMATIC, 2023) to illustrate gene flow between chromosomes, plasmids, and MGEs.

**Specialized Metabolism**

Secondary metabolite biosynthetic gene clusters (BGCs) were identified with antiSMASH v7.0 (Blin et al., 2023). Similarity networks were constructed using BiG-SCAPE v2.0 (Navarro-Muñoz et al., 2020), referencing MIBiG v3.1 (Medema et al., 2019), and visualized in Cytoscape v3.9.1 (Shannon et al., 2003).

**References**

Alcock, B.P., Raphenya, A.R., Lau, T.T.Y., Tsang, K.K., Bouchard, M., Edalatmand, A., Huynh, W., Nguyen, A.V., Cheng, A.A., Liu, S., Min, S.Y., Mistry, K., Niu, Y.D., Oloni, M., Speicher, D.J., Florescu, A., Singh, B., Tam, K., Tan, B., Thomas, M., To, D., Tong, Z., Whitney, H., Yeung, A., Zubyk, H.L., Pawlowski, A.C., Johnson, J., Brinkman, F.S.L., Wright, G.D., and McArthur, A.G. (2020). CARD 2020: antibiotic resistome surveillance with the Comprehensive Antibiotic Resistance Database. *Nucleic Acids Res.* 48(D1), D517–D525. doi:10.1093/nar/gkz935

Andrews, S. (2010). FastQC: a quality control tool for high throughput sequence data.

Bankevich, A., Nurk, S., Antipov, D., Gurevich, A.A., Dvorkin, M., Kulikov, A.S., Lesin, V.M., Nikolenko, S.I., Pham, S., Prjibelski, A.D., Pyshkin, A.V., Sirotkin, A.V., Vyahhi, N., Tesler, G., Alekseyev, M.A., and Pevzner, P.A. (2012). SPAdes: a new genome assembly algorithm and its applications to single-cell sequencing. *J. Comput. Biol.* 19(5), 455–477. doi:10.1089/cmb.2012.0021

Blin, K., Shaw, S., Kautsar, S.A., Medema, M.H., and Weber, T. (2023). antiSMASH 7.0: new features for the genomic mining of biosynthetic gene clusters. *Nucleic Acids Res.* 51(W1), W46–W50. doi:10.1093/nar/gkad343

Bolger, A.M., Lohse, M., and Usadel, B. (2014). Trimmomatic: a flexible trimmer for Illumina sequence data. *Bioinformatics* 30(15), 2114–2120. doi:10.1093/bioinformatics/btu170

Letunic, I., and Bork, P. (2021). Interactive Tree Of Life (iTOL) v6: an online tool for phylogenetic tree display and annotation. *Nucleic Acids Res.* 49(W1), W293–W296. doi:10.1093/nar/gkab301

Medema, M.H., Kottmann, R., Yilmaz, P., Cummings, M., Biggins, J.B., Blin, K., de Bruijn, I., Chooi, Y.H., Claesen, J., Coates, R.C., Cruz-Morales, P., Duddela, S., Düsterhus, S., Edwards, D.J., Fewer, D.P., Garg, N., Geiger, C., Gomez-Escribano, J.P., Greule, A., Hadjithomas, M., Haines, A.S., Helfrich, E.J.N., Hillwig, M.L., Ishida, K., Jones, A.C., Jones, C.S., Jungmann, K., Kegler, C., Kim, H.U., Kittelmann, M., Krug, D., Masschelein, J., Melnik, A.V., Mantovani, S.M., Monroe, E.A., Moore, M., Moss, N., Nützmann, H.W., Pan, G., Pati, A., Petras, D., Reen, F.J., Rosconi, F., Rui, Z., Sardar, D., Scherf, T., Schiewe, H., Skinnider, M.A., Soldatou, S., Sørensen, D., Steinbeck, C., Stroe, M.C., Tahlan, K., Tang, X., Tsueng, G., Varga, J., Vingataramin, M., Winkler, A., Yan, X., Yim, G., Yu, D., Karpinets,

T., Ziemert, N., and Glöckner, F.O. (2019). Minimum Information about a Biosynthetic Gene cluster (MIBiG) 2.0. *Nucleic Acids Res.* 47(D1), D625–D630. doi:10.1093/nar/gky940

Minh, B.Q., Schmidt, H.A., Chernomor, O., Schrempf, D., Woodhams, M.D., von Haeseler, A., and Lanfear, R. (2020). IQ-TREE 2: new models and efficient methods for phylogenetic inference in the genomic era. *Mol. Biol. Evol.* 37(5), 1530–1534. doi:10.1093/molbev/msaa015

Navarro-Muñoz, J.C., Selem-Mojica, N., Mullowney, M.W., Kautsar, S.A., Tryon, J.H., Parkinson, E.I., De Los Santos, E.L.C., Yeong, M., Cruz-Morales, P., Abubucker, S., Roeters, A., Lokhorst, W., Fernandez-Gutierrez, M., Kettleborough, C., Hadjithomas, M., Balunas, M.J., Bouslimani, A., Gallardo-Moreno, A.M., Pevzner, P., and Medema, M.H. (2020). A computational framework to explore large-scale biosynthetic diversity. *Nat. Chem. Biol.* 16, 60–68. doi:10.1038/s41589-019-0400-9

Page, A.J., Cummins, C.A., Hunt, M., Wong, V.K., Reuter, S., Holden, M.T.G., Fookes, M., Falush, D., Keane, T., and Parkhill, J. (2015). Roary: rapid large-scale prokaryote pan genome analysis. *Bioinformatics* 31(22), 3691–3693. doi:10.1093/bioinformatics/btv421

Panunzi, L.G. (2020). Corrigendum: sraX: a novel comprehensive resistome analysis tool. *Front. Microbiol.* 11:594635. doi:10.3389/fmicb.2020.594635

SankeyMATIC. (2023). A Sankey diagram builder for the web. Available online at: https://sankeymatic.com

Sayers, S., Usié, A., Rey, S., Glusman, G., Istrail, S., and Robinson, J. (2019). VICTORS: a web-based knowledge base of virulence factors. *Nucleic Acids Res.* 47(D1), D693–D700. doi:10.1093/nar/gky999

Seemann, T. (2014). Prokka: rapid prokaryotic genome annotation. *Bioinformatics* 30(14), 2068–2069. doi:10.1093/bioinformatics/btu153

Shannon, P., Markiel, A., Ozier, O., Baliga, N.S., Wang, J.T., Ramage, D., Amin, N., Schwikowski, B., and Ideker, T. (2003). Cytoscape: a software environment for integrated models of biomolecular interaction networks. *Genome Res.* 13(11), 2498–2504. doi:10.1101/gr.1239303

Wang, M., Goh, Y.X., Tai, C., Wang, H., Deng, Z., and Ou, H.Y. (2022). VRprofile2: detection of antibiotic resistance-associated mobilome in bacterial pathogens. *Nucleic Acids Res.* 50(W1), W768–W773. doi:10.1093/nar/gkac321
